# Trans-Driver: a deep learning approach for cancer driver gene discovery with multi-omics data

**DOI:** 10.1101/2022.06.07.495072

**Authors:** Hai Yang, Lei Zhang, Dan Zhou, Dongdong Li, Jing Zhang, Zhe Wang

## Abstract

Driver genes play a crucial role in the growth of cancer cells. Accurate identification of cancer driver genes is helping to strengthen the understanding of cancer pathogenesis and is conducive to the development of cancer treatment and drug-targe driver genes. However, due to the diversity and complexity of the multi-omics data, it is still challenging to identify cancer drivers.In this study, we propose Trans-Driver, a deep supervised learning method with a novel transformer network, which integrates multi-omics data to learn the differences and associations between different omics data for cancer drivers’ discovery. Compared with other state-of-the-art driver gene identification methods, Trans-Driver has achieved excellent performance on TCGA and CGC data Machine learning for multi-omics data integration in cancer. Among 20,000 protein-coding genes, Trans-Driver reported 185 candidate driver genes, of which 103 genes (about 55%) were included in the gold standard CGC data set. Finally, we analyzed the contribution of each feature to the identification of driver genes. We found that the integration of multi-omics data can improve the performance of our method compared with using only somatic mutation data. Through detailed analysis, we found that the candidate drivers are clinically meaningful, proving the practicability of Trans-Driver.

**Author summary:** Many methods have been developed to identify cancer driver genes. However, most of these methods use single-omics data for cancer driver gene identification. Multi-omics-based methods for cancer driver gene identification are rare. Trans-Driver uses deep learning to process multi-omics data and learn the relationships between multi-omics data for cancer driver gene prediction. We have predicted 185 candidate cancer driver genes out of among 20,000 protein-coding genes. Also, we performed cancer driver gene prediction on 33 cancer types, and we identified the cancer driver genes corresponding to each cancer type. And, we observed that the predicted cancer driver genes were shown to have a role in cancer progression in recent studies. Our proposed method for cancer driver gene identification using multi-omics data has improved performance compared to using mutation data alone.

## Introduction

Cancer is a genomic disease that poses a severe threat to human life and health [1]and is a significant cause of disease-related morbidity and mortality worldwide [2]. It is an abnormal and uncontrolled disease of cells caused by genetic mutations [3, 4]. Previous studies have shown that cancer cells arise, develop, metastasize, and worsen due to the accumulation of severe mutations in the genome of the body’s matching tissue cells over time [5, 6]. Driver mutations disrupt normal cell growth and promote cancer development, while most passenger mutations do not contribute to cancer development. Genes with the driver mutations are cancer driver genes [7]. The discovery of cancer driver genes is critical for cancer diagnosis, prevention, clinical treatment, and development of cancer-targeted drugs [8, 9]. With the development of next-generation sequencing technologies, some international cancer research projects, such as The Cancer Genome Atlas (TCGA) [10] and International Cancer Genome Consortium (ICGC) [11], have produced many omics data (such as Somatic mutation, DNA methylation, copy number), which contributed to the study of cancer driver genes [12].

Over the past decade, numerous approaches have been developed to identify driver genes. Most of these methods (such as MuSiC [13], ActiveDriver [14], OncodriverFML [15], VEST [16]) are based on statistical models that use somatic mutation data to estimate background mutation rates combined with mutation rates of genes to identify driver genes. MuSiC used statistical methods to decide mutational salience in cancer, distinguishing significant mutations as driver mutations among passenger mutations. ActiveDriver identified the signaling regions of substantial mutations in proteins. This method complemented the standard frequency-based mutation significance approach and helped explain rare but site-specific mutations. OncodriverFML introduced a local mutation background model to calculate the mutation bias of genomic for the analysis of coding and non-coding somatic mutation patterns in genomic regions to identify driver mutations. Researchers have used machine learning methods to improve cancer drivers’ identification prediction accuracy. VEST was a machine learning-based supervised classification method for classing missense mutations that alter protein to identify possible missense mutations. Some methods integrate somatic mutations and other biological knowledge to improve identification accuracy. Tokheim et al. proposed the 2020plus, a Random Forest-based machine learning prediction algorithm for predicting oncogenes and tumor suppressor genes in somatic mutations, using features such as capture mutation clustering, evolutionary conservation, prediction of variants, mutation, gene network connectivity, and other covariates [17]. The e-Driver algorithm identified cancer driver genes by the intrinsic distribution pattern of somatic missense mutations among functional regions of proteins [13]. CHASM used conservation scores and protein structure features to predict the driver mutations most likely to produce changes and promote cancer cell spread [18]. Bailey et al. developed the CompositeDriver method, which considered the effects of mutation recurrence and the computational mutation method for identifying positive selection signals [7].The PNC algorithm identifies the personalized driver genes by employing the structure-based network control principle on the genetic data of individual patients [19]. For all mutations within the protein-coding region of a gene, a composite score was calculated by multiplying mutation recurrence by the impact score [20].

Integrating multi-omics data is essential in cancer research [21–23]. Several recent studies have shown that multi-omics data integration is more helpful in exploring cancer driver genes compared to using only somatic mutation data [24–26]. Liu et al. developed an online cancer driver gene database, DriverDBv3, using various published methods through multi-omics data to improve the study of comprehensive cancer omics data by identifying driver genes [27]. Singh et al.used simulations and benchmark multi-omics to identify known and novel multi-component biomarkers between multiple phenotypic groups by selecting common features from multi-omics data sets [28]. deepDriver is a proposed method for cancer driver gene identification by convolving mutation-based features of genes and their neighbors in a similarity network [29].LOTUS is a machine-learning-based approach that allows for the integration of various types of data versatile, including information about gene mutations and protein-protein interactions [30]. Our previous work developed a non-parametric Bayesian framework based on multivariate statistical modeling: iDriver, which integrated somatic mutations, gene expression, DNA copy number, DNA methylation, and protein plenty provided by The Cancer Genome Atlas (TCGA) to identify cancer driver genes [31].

Although many computational approaches have emerged to use the next-generation gene sequencing technologies for cancer drivers’ discovery, the goal of achieving a complete cancer drivers catalog remains challenging. Many methods have significant gaps in prediction consistency and the number of predicted cancer driver genes [17]. Significant challenges still go with this task. In recent years, with the booming field of artificial intelligence (AI), deep learning has increasingly overlapped with cancer medicine [32]. The vast amount of available biomedical data filtered through the analytical power of AI promises to revolutionize cancer research, diagnosis, and care [33]. Deep learning can predict genomic changes based on morphological features learned from digital histopathology [34]. Deep learning has been used to characterize multi-omics data, molecular subtyping of tumors, and predict patients’ survival rates [35–38].

Since multi-omics data integration can effectively identify cancer driver genes and deep learning has advantages for multi-layer abstract representation of many complex data to explore novel cancer drivers better. This study proposes a deep supervised learning approach, Transformer-Driver (Trans-Driver). We innovatively proposed a multi-layer perceptron-Transformer network with multi-omics data integration to deal with the imbalance of driver and passenger genes in the training set (Fig 1). We compared Trans-Driver with other driver gene discovery methods such as MuSiC, ActiveDriver, e-Driver, CompositeDriver, CHASM, and 2020plus. Our approach achieved outstanding performance on the two benchmarks data sets (TCGA drivers set and COSMIC CGC set). Among 20,000 protein-coding genes, Trans-Driver predicted 185 alternative driver genes, of which 103 genes (55%) were included in the gold standard CGC data set. Finally, we analyzed the contribution of each feature to identifying cancer driver genes, and the results suggested that mutation data contribute most to the identification of cancer driver genes. However, methylation data are also essential for identifying driver genes in some cancers.

**Fig 1.**
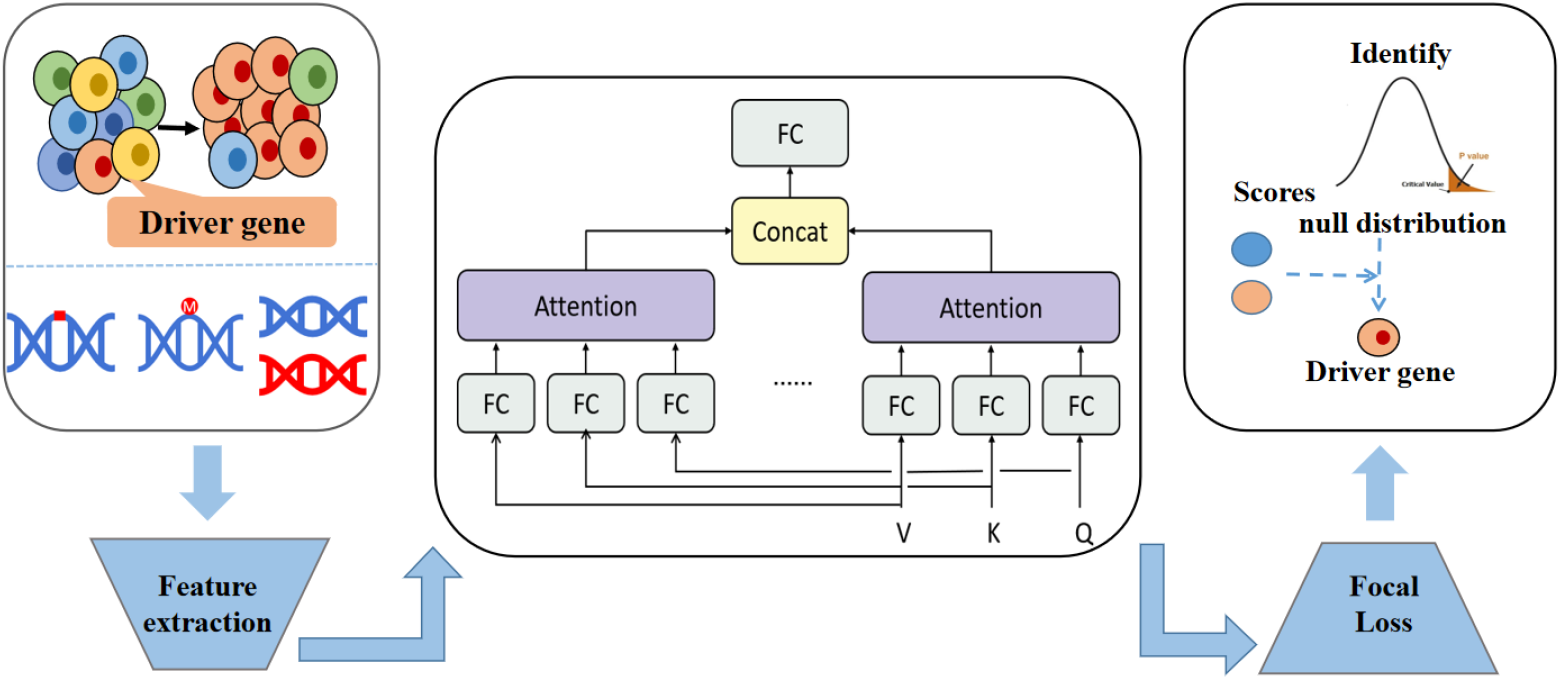
Overview of the Trans-Driver framework. Identification of cancer driver genes by integration of multi-omics data using the transformer structure.We first collected and processed multi-omics data, then used the Transformer model to learn features, scored the model using Focal Loss, and obtained the final cancer driver genes based on the constructed background distribution.

## Materials and methods

### Overview of Trans-Driver

The input of Trans-Driver is the multi-omics data across 33 TCGA tumor sets, and the output of the approach is the probability that the gene is the cancer driver. The framework can be divided into two parts. The first part extracts the features from the multi-omics data. The second part, the multi-layer perceptron-Transformer network [39], is developed to learn the features’ representations and calculate the driver gene scores (see Supplementary Note 1 for details).

### Data preprocessing and feature selection

This study’s cancer driver gene discovery is based on the TCGA data set [7]. We collected the DNA methylation, Copy number variation (CNV) data, and somatic mutation data across 33 types of cancer from the TCGA Pan-cancer Atlas [40](See Supplementary Note 2 for a detailed description of the data).

We selected the ‘Variant_Classification’ (‘Frame_Shift_Del’, ‘Frame_Shift_Ins’, ‘In_Frame_Del’,’In_Frame_Ins’,’Missense_Mutation’,’Nonsense_Mutation’,’Silent’, ‘Non-stop_Mutation’,’Splice_Site’, ‘Translation_Start_Site’) and ‘Variant Type’(‘DEL’, ‘INS’, ‘SNP’) columns from the somatic mutation spectrum to extract somatic mutation data features. We counted the number of various somatic mutations corresponding to each gene, defaulting to 0 when a particular type of mutation did not occur at all. Finally, we generated mutation-related features (28 values) with our feature extraction module.

To extract the features of the CNV data, we obtained the matched data for all the samples based on the ‘Tumor_Sample_Barcode’ column in the mutation profiles and then calculated the mean value of the CNVs. Afterward, we obtained the corresponding relationship between ‘Hugo_Symbol’ and ‘Tumor_Sample_Barcode’ in somatic mutation data. We got the corresponding relationship between protein-coding genes and copy number variation intervals and then calculated the mean and variance of CNVs of each gene. To consider both upregulation and downregulation of copy number variation, we counted the mean and variance of the two cases separately as the copy number variation features (6 values). Since the CNV-related features are continuous values, we fill in the missing data by calculating the mean of the missing values.

We obtained DNA methylation data from the TCGA Pan-cancer Atlas. Similar to the CNV data processing, the mean and variance of each gene were calculated (2 values), and the DNA methylation data were mapped to specific genes based on the CpG sites (CG) islands and the linked genes. Finally, we also introduced some other features, including the gene degree (the degree of the gene in a general protein-protein interaction (PPI) network), the centrality between genes, the DNA replication time of the genes, and the chromatin status of the genes. We produced the features (46 values) based on multi-omics data as model inputs for Trans-Driver.

### The deep learning model of Trans-Driver

The differences and associations between the omics data during the model’s training process must be considered. The advantage of deep learning models is the accurate modeling of complex inputs and the ability to consider the differences and connections between different types of omics data through the attention mechanism. The neural network of Trans-Driver is divided into three modules (Fig 2): perceptron, transformer with multi-headed attention block, and the classifier. The perceptron module extracts the omics data features using a linear layer. The transformer module learns the relationships between the features extracted by the perceptron module. The extracted low-dimensional representations of the input are fed to the classifier module for gene classification. Let *x*_*k*_ be the input data of the perceptron network. The input signal is linearly transformed to become the output signal of the perceptron model. To mitigate the occurrence of overfitting, a dropout layer is added to the perceptron module, which first uses the Bernoulli function to generate a vector with probability *p* randomly:

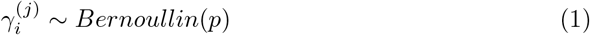

where *x*_*ki*_ is the input of the first node of the first set of data, *v* is the output of the *j* th node of the k th set of data, *w*_*ij*_ is the weight,*b*_*kj*_ is the deviation value, and *f* () is the activation function. We choose the Gaussian Error Linear Unit (GELU) function [41] as the nonlinear activation function of the network:

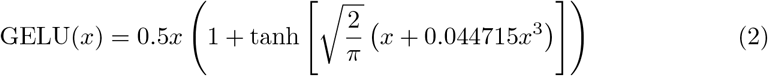

next, we introduce the self-attention mechanism, an attention method to improve the accuracy of local feature representation by aligning internal and external information observation. In this mechanism, the input signal is divided into query unit query (*q*), key-value unit key (*k*), and representation unit value (*v*), and *q* and *v* are in one-to-one correspondence. The attention calculation formula is as follows:

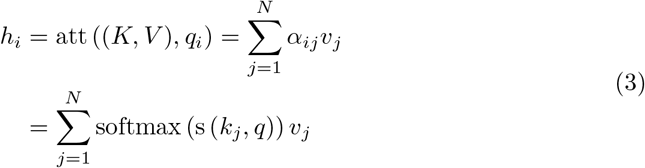

where *att*() is the attention distribution, *s*() is the attention scoring function, and *softmax*() is the normalization function. We use the scaled dot product as the attention scoring function: firstly, calculate the dot product of *x*_*i*_ and *q*, secondly, divide by 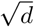, *d* is the dimensional value of *x*_*i*_. The weight distribution on *v* is obtained by softmax, and finally, the weighted value of *v* is obtained by the dot product. In practice, a parallel estimate is performed using batch processing, so the matrix representation of the attention mechanism takes the following form:

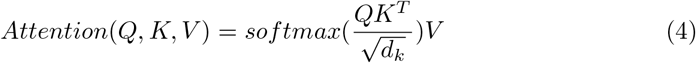

where *Q = W*^*q*^*y, K = W*^*k*^*y, V = W*^*v*^*y, d*_*k*_ is the number of columns of the *Q,K* matrix. After getting *QK*^*T*^, the attention coefficients are obtained using the softmax function. *W*^*q*^,*W*^*k*^, and *W*^*v*^, are the three different weight parameter matrices to be trained by the model, and *y* is the output data of the previous perceptron module. Next, we stitched the results of the self-attentive mechanism for *h* times and performed another linear transformation yielded the results with the following equation:

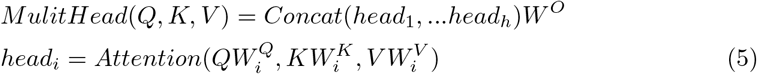

where 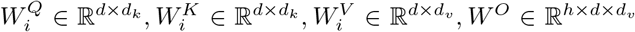. d_model means the output dimension of the perceptron module, *h* = 2,*d*_*k*_ = 4,*d*_*v*_ = 8. The Multi-Head Attention block contains multiple Self-Attention layers, firstly, the input *y* is passed to each of the *h* different Self-Attentions, and the different outputs *Z*_*i*_ are computed. Then they are stitched together to get the final output.

**Fig 2.**
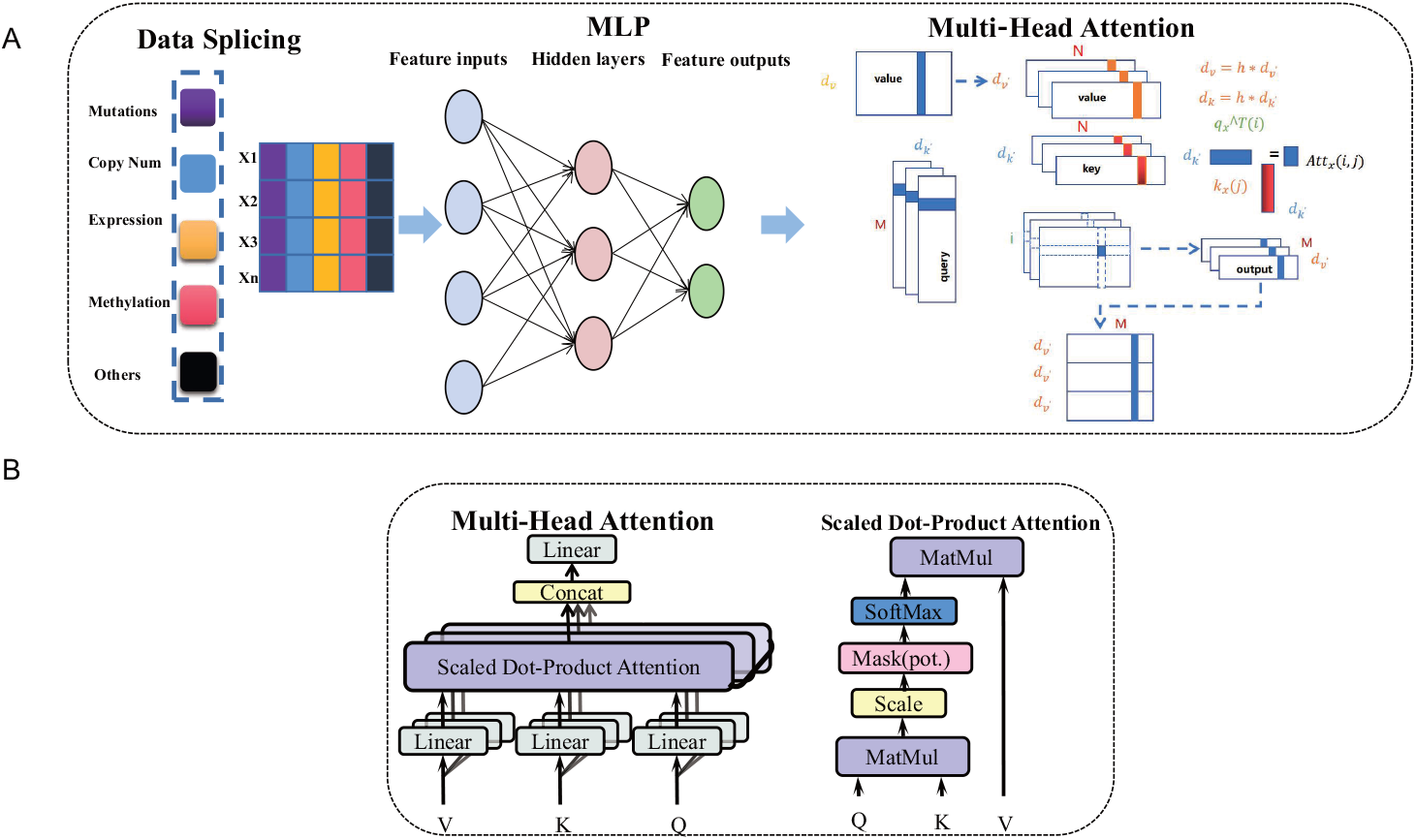
The deep learning model of Trans-Driver. (A)The network of the model contains data stitching, MLP feature extraction, and transformer and scaled dot product attention is used in the transformer block. We first processed the multi-omics data, then performed a linear transformation, and then processed the data using a multi-headed attention block.(B)Concrete implementation of the multi-headed attention block.

The results obtained by the multi-headed attention block are transformed by the linear layer and then by the activation function to get the results. We use the Focal loss function [42] as the loss function to deal with the imbalance of positive and negative samples in the training process:

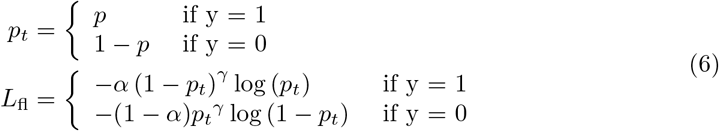

where *γ* and *α* are the model’s hyperparameters, and all network parameters are updated synchronously to train the model until the gradient descent converges. *γ* reduced the loss of easily classified samples, making the model focus on the problematic and misclassified samples, and fully exploring the challenging examples. Since the number of drivers is smaller than the number of passengers, *α* can be set to make the model focus more on the driver genes. We explored the values of alpha and gamma parameters(Fig S1) and finally selected *α* = 0.4 and *γ* = 1.5.(see Supplementary Note 3 for details, The loss function of the model is shown in Fig S2).

### Construction of the background distribution

We constructed the empirical background distribution with the Monte Carlo simulation [17]. Specifically, single nucleotide variants (SNVs) are moved randomly to new matched positions in the genome for each gene, keeping the total number of SNVs fixed. The matched positions must have the same base context as the original positions, such as C*pG, CpG*, TpC*, G*pA, A, C, G, and T. The number of SNVs remains constant, but the mutation outcome may change. We put the gene feature data from the simulation into the Trans-driver model for scoring and formed an empirical null distribution based on the scoring. The number of simulations was set to 100,000. After the above Monte Carlo simulations produced the empirical null distribution, the score was obtained using real data in the Trans-driver model for each gene. To calculate the p-value of the gene, we use the score of the simulated gene with a score greater than or equal to the threshold. Finally, the Benjamini-Hochberg method was used to control for error incidence to calculate the q-value of the gene, and we considered a gene to be a driver if its q-value was less than 0.05. To estimate the background distribution of driver gene scoring on the 33 types of tumors more accurately, we performed simulations of mutations for each cancer type.

### Data set and evaluation metrics

We selected the 71 oncogenes and 53 tumor suppressor genes [17]to train the neural network as the positive samples. We resampled the same number of genes among the remaining genes as the negative samples. The training samples were shuffled to generate the model’s training set, which contained 248 genes.

We constructed a benchmark data set to confirm the model’s performance in discovering cancer drivers. We used the 299 driver genes reported by TCGA [7]as the positive samples, and negative samples were randomly selected from other coding regions genes.We used different data sets for training and testing to avoid the potential over-fitting problem. We executed deduplication on the test set to remove the positive and negative sample genes used in the training set and got the TCGA benchmark data set with 380 genes.

We chose a more extensive data set to test the model. We downloaded the COSMIC CGC data [43] (v94) and selected 680 cancer genes with somatic mutations. The negative sample selection and deduplication were performed, and we finally got the CGC test set with 1058 genes.

In this study, the Receiver Operating Characteristic (ROC) matric was used to evaluate the method, and the ROC curves were used to compare the false positive rate (FPR) to the true positive rate (TPR) at different thresholds. The false-positive rate represents the ratio of false positives to false positives and true negatives, and the true positive rate represents the ratio of true positives to true positives and false negatives. True-positive shows that the model successfully identified the initially driver gene, but false-positive shows that the model incorrectly identified the original passenger gene. True negative shows that the model successfully identified the original passenger gene as a passenger gene. False-negative shows that the model predicted the original driver gene as a passenger gene. AUROC shows the area under the ROC curve. We performed enrichment analysis on the CGC data set and TCGA data set using Fisher’s test to better evaluate the truth of driver genes predicted by each method.

To analyze more deeply which omics data are most relevant to cancer driver genes, we assessed the feature importance of the integrated multi-omics data using a Random Forest (RF) approach by analyzing the contribution of each feature to each tree in the Random Forest. Here we used the Gini index calculation by taking the average value and then comparing the importance of the contribution between features:

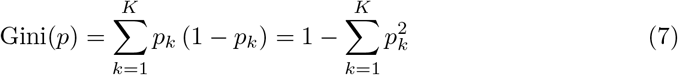

where *K* is the *K* categories and *P*_*k*_ is the sample weights of category.

## Results

### Pan-cancer driver genes discovery with Trans-Driver

To confirm the effectiveness of the proposed Trans-Driver algorithm in identifying cancer driver genes, we first evaluated the results of Trans-Driver on the Pan-cancer data set. We constructed two benchmark data sets: the driver gene set published by TCGA and the COSMIC CGC gene set. We used six state-of-the-art driver gene identification methods: ActiveDriver, e-Driver, MuSiC, CompositeDriver, CHASM, and 2020plus for the performance comparison. The driver gene scores of the approaches were obtained from the previous study [7]. We AUROC as a metric for performance evaluation between the different algorithms on both benchmark data sets. We observe from (Fig 3A) the Trans-Driver algorithm achieves the best performance on the TCGA data set (AUROC=0.911), followed by 2020plus (AUROC=0.900), MuSiC (AUROC=0.874), CompositeDriver (AUROC=0.839), e-Driver (AUROC=0.753), ActiveDriver (AUROC=0.641), and CHASM (AUROC=0.624). In Fig S3A, We also calculated the AUPRC values of each methods Trans-Driver (AUPRC=0.904) and calculated the Mann-Whitney significant difference check to obtain p-values (p-value= 2.540e-43), followed by 2020plus (AUPRC=0.918, p-value=6.994e-41), MuSiC (AUPRC=0.852, p-value=2.250e-36), CompositeDriver (AUPRC=0.852, p-value=2.51e-29), e-Driver (AUPRC=0.767, p-value 7.707e-17), ActiveDriver (AUPRC=0.716, p-value=5.934e-06), and CHASM (AUPRC=0.615, p-value=3.517e-05).

**Fig 3.**
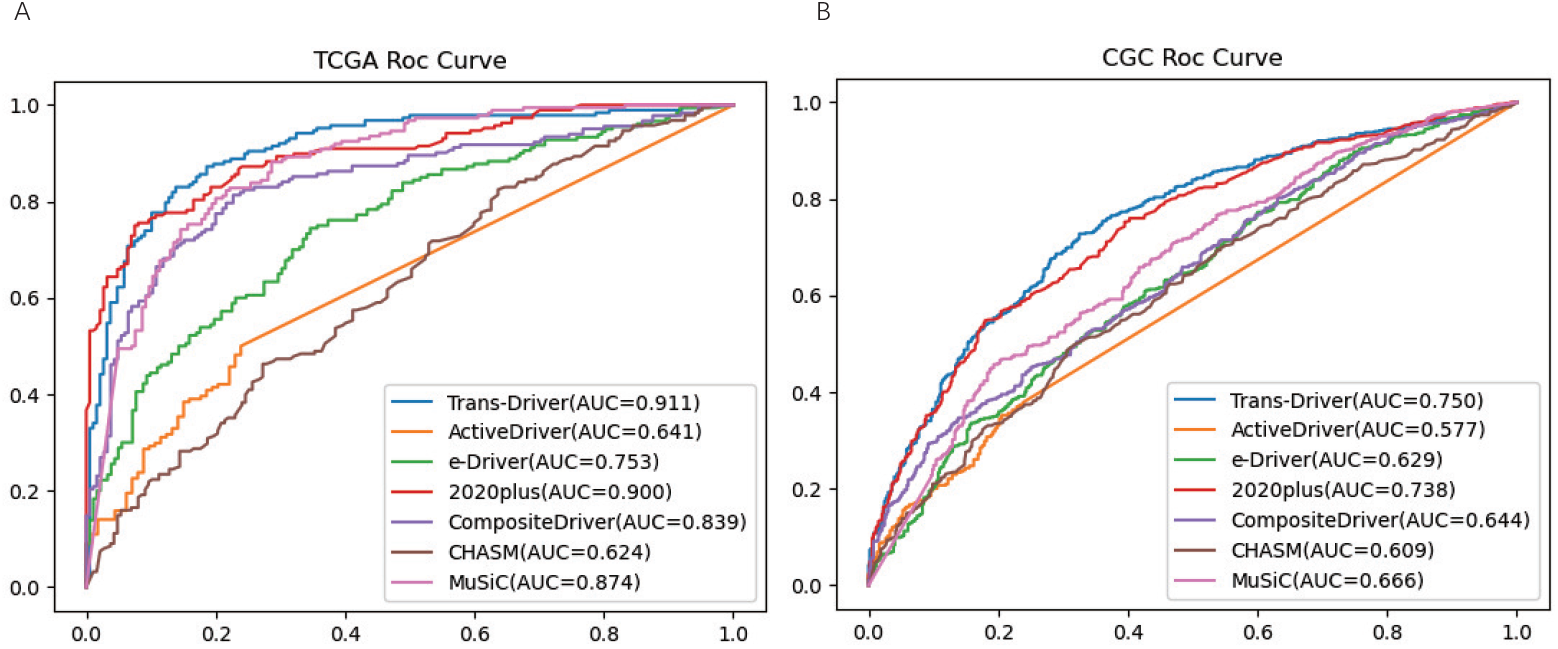
The performance evaluation of different cancer driver gene prediction methods. Trans-Driver and other methods include ActiveDriver, e-Driver, 2020plus, CompositeDriver, CHASM and MuSiC. (A) ROC curves of various methods on TCGA data set, (B) ROC curves of various methods on CGC gene set.

Next, we compared the performance of Trans-Driver and the other approaches on the COSMIC CGC data set. Since the genes in CGC are from different cancer types than the 33 cancers of interest in this analysis, we can see from Fig 3B and in Fig S3B that most of the algorithms’ performances were decreased. Nevertheless, the Trans-Driver algorithm also outperforms the other methods on CGC (AUROC=0.750, AUPRC=0.751, p-value=4.864e-45), followed by 2020plus (AUROC=0.738, AUPRC=0.740, p-value= 5.602e-41), MuSiC (AUROC=0.666, AUPRC=0.644, p-value=1.014e-20), CompositeDriver (AUROC=0.644, AUPRC=0.661, p-value=7.916e-16), e-Driver (AUROC=0.629, AUPRC=0.625, p-value=1.677e-12), CHASM (AUROC=0.609, AUPRC=0.611, p-value=7.585e-10), and ActiveDriver (AUROC=0.577, AUPRC=0.669, p-value=1.653e-05). In general, Trans-Driver achieves a stable performance improvement on both gold standards.

The AUROC metric reflects the overall performances of each method on all 20,000 coding region genes. Still, it is also important to assess the significance of enrichment of top-ranked genes with the gold standard. For this purpose, after simulating the empirical zero probability distribution through Monte Carlo, we use Fisher’s exact test to calculate the empirical p-value. Simultaneously, we performed FDR correction for p-values of all genes to get q-values. We considered Trans-Driver on the Pan-cancer data set with q-value *<*0.05 as the driver genes (185 genes). For other methods, we use the q-value cutoff in the previous study ([7]) to select their top-ranked genes: CompositeDriver (78 genes, q-value=0.05), ActiveDriver (140 genes, q-value=0.0001), OncodriveCLUST (145 genes, q-value=0.05), 2020plus (164 genes, q-value=0.05),e-Driver (233 genes, q-value=0.1), MuSiC (2923 genes, q-value=1e-10), CHASM (2933 genes, q-value=0.1).The 185 driver genes reported by Trans-Driver on Pan-cancer are the most enriched with both the TCGA gene set and the CGC gene set. In summary, the enrichment of Trans-Driver both on TCGA and CGC data sets is better than other comparative algorithms, showing the genes predicted by our Trans-Driver algorithm for Top N are equally reliable.

Genes identified by multiple methods are more likely to be actual cancer driver genes [17]. We counted the over-lap rate of driver genes predicted by each method on Pan-cancer and those pinpointed by other methods (Fig 4A) to show the consistency in predicting driver genes. We divide the cancer driver genes predicted by each method into four groups, the first group being cancer driver genes specific to each method predicted, the second and third groups being cancer driver genes predicted by 2 and 3 methods, respectively, and the last category being driver genes predicted by at least four methods. From Fig 4A, we found that the driver genes predicted by Trans-Driver have a high agreement with other methods (The proportion of genes predicted that were also predicted by other methods was at least 92.97%). We also found that Trans-Driver also reported some novel driver genes (7.03%) due to deep learning methods.

**Fig 4.**
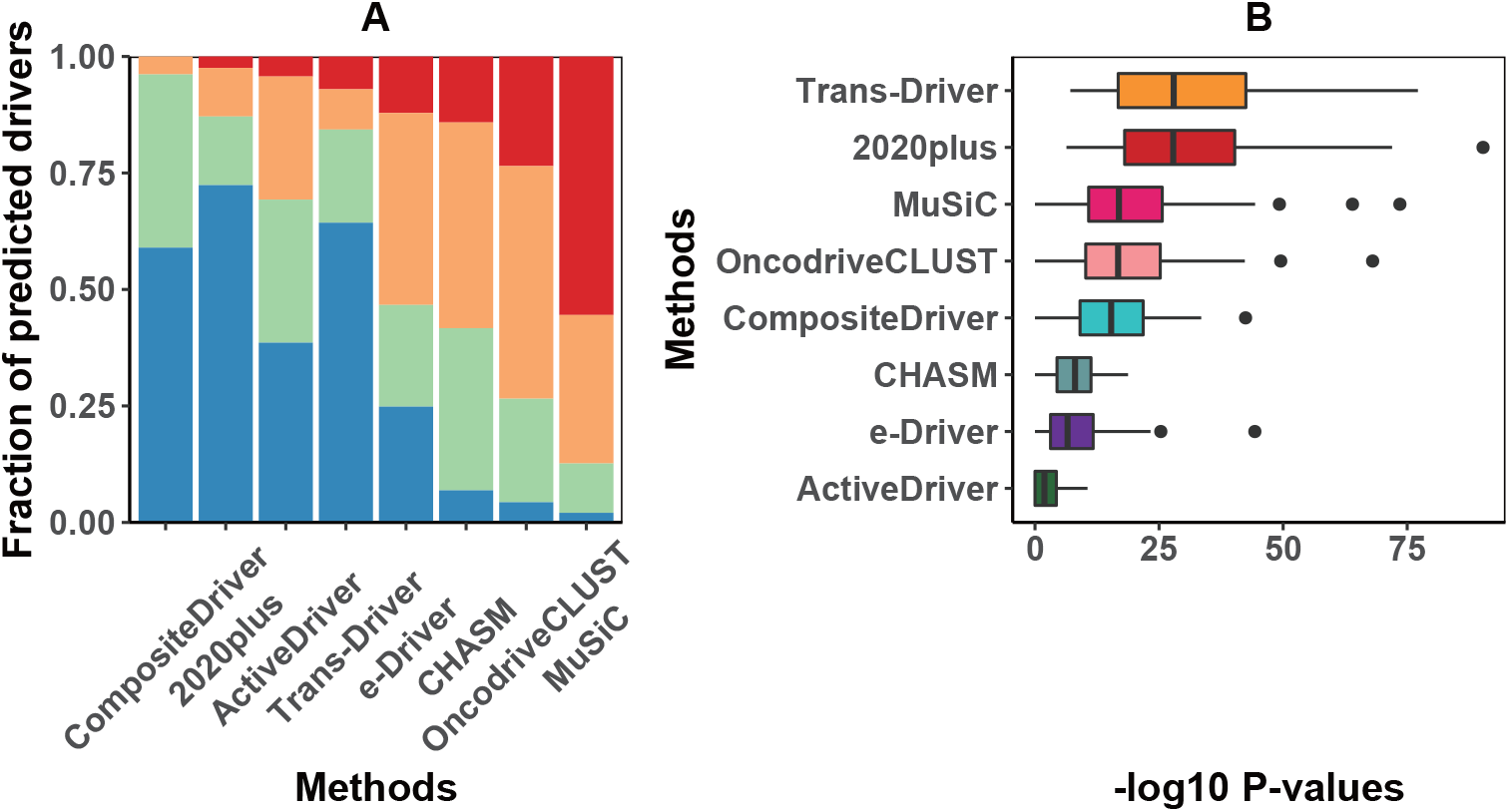
The analysis results of the methods: (A) The proportional consistency of the predicted driver genes with the eight methods. (B) Enrichment levels of eight methods on 33 cancer types from TCGA data.

We also compared the Trans-Driver with the traditional Multilayer Perceptron (MLP) model. From Fig S4, we can see that our proposed model outperforms the MLP model in both pr and roc on both the TCGA and CGC data sets.

### Analysis of the performance of Trans-Driver and other identification algorithms on 33 TCGA cancer data sets

Since we have observed that Trans-Driver has superior performance on the Pan-cancer data set. To study the driver genes on each cancer type (especially some challenging rare tumor types), we apply Trans-Driver on 33 cancers publicly available to the TCGA data set for driver gene discovery. By integrating the matching four types of omics data on different tumor types and modeling them with Trans-Driver, we finally succeed in predicting a reasonable number of candidate driver genes on each tumor set. Trans-Driver predicted the highest number of driver genes on UCEC tumors (147 genes) while predicting the lowest number of driver genes on KICH tumors (6 genes). The candidate drivers of Trans-Driver across each tumor type are reported in Table S1.

To justify the driver gene prediction results of Trans-Driver on each cancer type, we compare the performance of Trans-Driver with other approaches on all cancer types. Since there are fewer data for each cancer type than the Pan-cancer set and few known positive samples, we only focus on the enrichment of different algorithms with known driver genes on each type of tumor (the Fisher test was used for the evaluation, and the -log10 p-value was used as the performance metric). We measured the enrichment of each method on the 33 cancer data sets with the TCGA gene set and compared the performance between different algorithms. We reported the Fisher’s exact test -log10 p-value of enrichment analysis of all methods on 33 cancer data sets (Fig 4B). The median of enrichment results of trans driver on 33 cancer types is the highest (median Fisher’s exact test -log10 p-value = 27.95). The median of enrichment results of other methods are 2020plus (27.85), followed by MuSiC (22.16), OncodriveCLUST (16.75), CompositeDriver (15.35), CHASM (8.07), e-Driver (6.56), and ActiveDriver (1.90). We also find that Trans-Driver achieves the best performance on 14 out of 33 cancer types. The enrichment results for e-Driver, CHASM, and MuSiC were 0 on the CHOL, while the enrichment result for Trans-Driver was 13.31 for the best result on the CHOL. The performance of each method for enrichment analysis across all cancer types is shown in Table S2. Due to the considerable variation of mutation rate in different cancer types (10.27 mutations per capita in PCPG cancer and 986.71 mutations per capita in UCEC cancer), it is difficult to obtain a reasonable number of driver genes in all cancer types. We observed that ActiveDriver and e-Driver could not correctly predict the driver genes in CHOL, KICH, and MESO (no intersection with the known driver gene set of TCGA corresponding cancer types). At the same time, Trans-Driver can obtain stable identification results in all 33 cancer data sets through the integration of multi-omics data. Four drivers were successfully identified on CHOL, two on KICH, and five on MESO. We also performed an enrichment analysis of two novel methods published in 2020 [44, 45]across 33 cancer types (see Table S3 for details), and the performance of Trans-Driver is still better than the two novel methods.

### Explore the potential driver genes predicted by Trans-Driver

We have elucidated that Trans-Driver successfully predicts a reasonable number of potential driver genes on all 33 cancer types and achieves excellent performance on TCGA gene sets for different cancer types. Next, we explored the candidate drivers reported by Trans-Driver on the Pan-cancer data set and on 33 cancer types. In the Pan-cancer data set, Trans-Driver identified 185 potential driver genes. Among them, 83 genes such as TP53, BRCA1, RB1, PTEN, and ATM (see Table S4 for details) are found in the TCGA and CGC gene sets, which are highly reliable cancer drivers. Meanwhile, LRP1B, FAT4, MUC16, CBFB, and other genes identified by Trans-Driver have been reported in the CGC data set, but they are tough to be found by computational methods.

Moreover, a literature study found that the following genes reported by Trans-Driver are likely to be potential driver genes. Among BLCA tumors, KMT2D exhibits the highest mutation rate of 26.9% in bladder cancer, and KMT2D knockdown inhibited bladder cancer cell viability, migration, and invasion in vitro. It may play a role in bladder cancer development [46]. Deletions of FOXA1 resulted in sexually dimorphic changes in uroepithelial differentiation, and loss of FOXA1 expression is associated with aggressive uroepithelial carcinoma of the bladder and increased tumor proliferation and invasion [47]. GPS2 is a multifunctional protein that plays an essential role in the inflammation and metabolism of adipose, liver, and immune cells.GPS2 was recently identified as a significantly mutated gene in breast cancer and other malignancies and proposed as a putative tumor suppressor [48]. Mutations in KMT2D methyltransferase are highly recurrent, and KMT2D mutations associated with diffuse large B-cell lymphoma impair KMT2D enzyme activity, and overall H3K4 methylation is reduced in diffuse large B-cell lymphoma cells. Early deletion of KMT2D, a tumor suppressor gene, acts by remodeling the epigenetic landscape of cancer precursor cells to promote lymphoma development [49]. In LUAD, SETD2 is mutated or loss-of-function in solid cancers, including renal, gastrointestinal, lung, pancreatic, and osteosarcoma. Mutation or loss-of-function of the SETD2 gene leads to dysfunction of the corresponding tumor tissue protein, resulting in tumorigenesis, progression, chemoresistance, and unfavorable prognosis, suggesting that SETD2 may act as a tumor suppressor [50]. CUL3 is a member of the ubiquitin ligase complex that functions in the oxidative stress response pathway. Evidence suggests that CUL3 acts as a tumor suppressor in non-small cell lung cancer [51].

In particular, we explored the novel driver genes reported by Trans-Driver but not in the CGC/TCGA data sets. On the Pan-cancer data set, Trans-Driver pinpointed 185 genes, of which 68 were not in TCGA and CGC gene sets. Among these, the SPTA1 gene was found to promote the development of GBM cancer [52]. CSMD1 prevents LIHC by suppressing cell invasion [53]. Moreover, Trans-Driver identified 505 potential driver genes across 33 cancers, of which a total of 413 genes are not in the CGC/TCGA data sets.Some of these novel drivers play a critical role in cancer development on BRCA, COAD, and other cancers. For example, On BRCA, Trans-Driver identified ELAVL1, an emerging target for breast cancer therapy, especially in metastatic breast cancer. ELAVL1 is a ubiquitously expressed post-transcriptional regulator, and cytoplasmic ELAVL1 accumulation is associated with poor overall and disease-free survival in high-grade malignancies. It can be an independent prognostic factor in poor clinical outcomes in breast cancer [54]. On the COAD data set, Trans-Driver predicts the PMG5 gene, which has been found to play an essential role in proliferation, invasion, and migration in COAD by recent studies, and decreased PGM5 is associated with poor prognosis [55] (See Supplementary Note 4 for details).

### Analysis of the contribution of different omics data to the identification results of Trans-Driver

The input of Trans-Driver includes somatic mutations, methylation data, CNVs, and other information such as gene length and conservation scores. To illustrate that the integration of multi-omics data can indeed improve the performance of the method (especially for algorithms based on supervised learning models such as Trans-Driver), first, we compare the performance of Trans-Driver using the integration of multi-omics data as features and using only mutation-related features under two test scenarios, Pan-cancer, and each cancer. We used Fisher’s exact test (-log p-value) on Pan-cancer and 33 cancer types, respectively, and the prediction performance (AUROC) on TCGA and CGC. We obtained the pairwise performance comparison plots for the five scenarios (Fig 5A). The performance of our approach with the multi-omics input on both test sets, TCGA and CGC, has improved somewhat over the model using only mutation-related features, with the value of AUROC improving by 1.33% on the TCGA gene set and 4.31% on the CGC test set. The enrichment performance of the model is also significantly improved in the enrichment analysis, both on Pan-cancer and on each cancer. The enrichment performance is enhanced by 6.02% using data from multi-omics data on TCGA compared to mutation-only data, and the enrichment result is enhanced by 16.04% on CGC. The enrichment result was improved by 25.90% on each cancer median. Under each test scenario, the performance of Trans-Driver has a stable and steady performance improvement compared with just mutation features by supervised learning of team multi-omics features, indicating the necessity of multi-omics integration.

**Fig 5.**
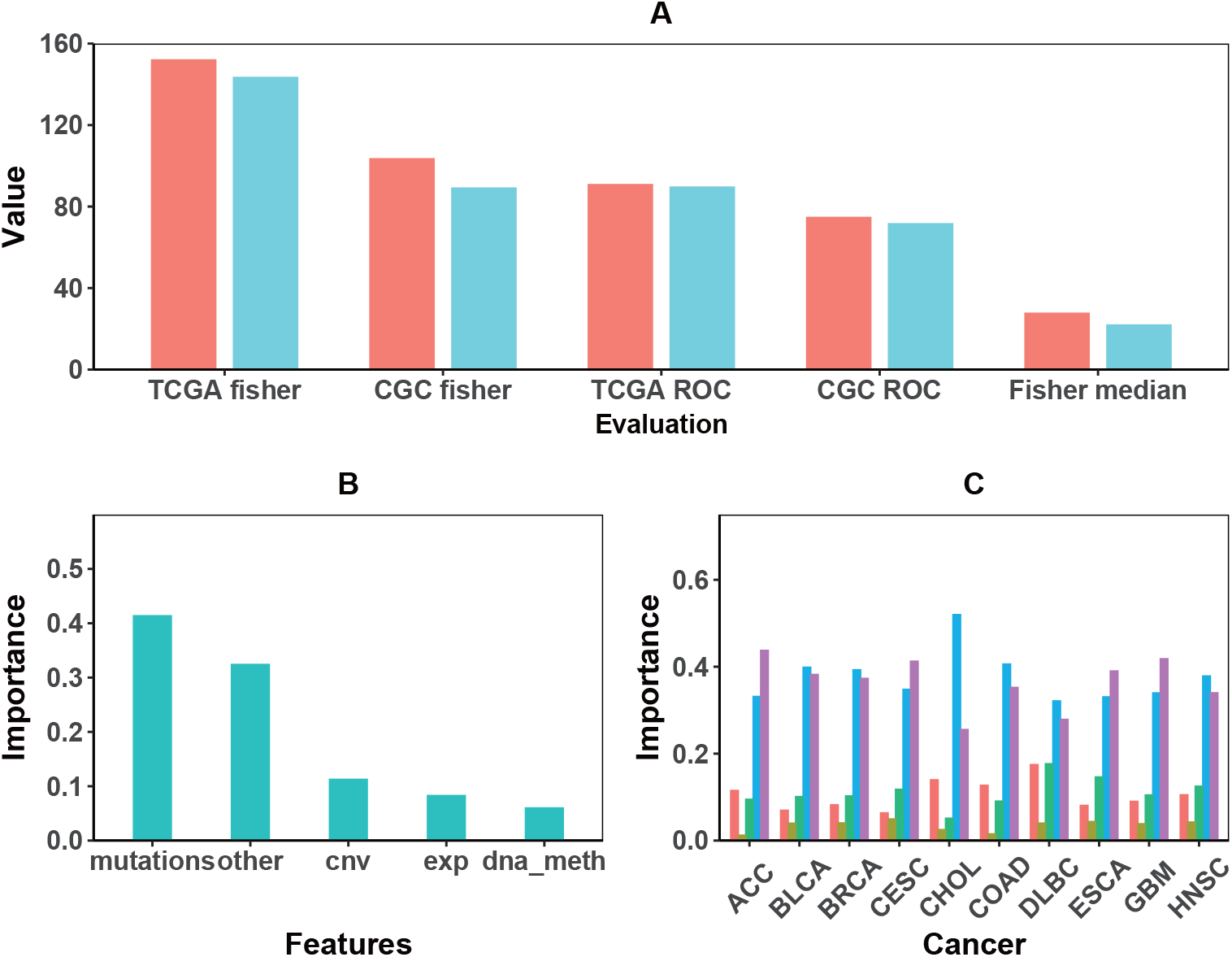
The feature importance analysis of multi-omics data: (A) Modeling with multi-omics features or mutation features was compared in the CGC data set and TCGA data set. (B)Analysis of the multi-omics features using random forest. (C)The proportion of multi-omics features contributes to the identification results.

To explore the importance of non-mutation data on the identification results of Trans-Driver, we calculated the contribution of these distinct omics data to the results of Trans-Driver. We introduced the Random Forest method to measure the input features’ influence on Trans-Driver’s identification results. The input of RF was the original integrated features and the identification scores of the Trans-Driver. The output of RF was the Gini importance scores of the features. Finally, we summed all the Gini importance scores belonging to each platform and quantified the contribution of different omics data to the final identification results. All the features are classified into five categories: mutation data, copy number, DNA methylation, expression, and others. Firstly, we calculated the Gini scores of each feature and averaged the scores of each category. The importance percentage of each feature category is shown in Fig 5B. We observed that mutation-related features contributed the most to identifying driver genes (41.51%). Still, other omics features are also important: other (32.54%), CNV (11.40%), gene expression (8.41%), and DNA methylation (6.14%). We also analyzed the contribution of features on ACC, BLCA, BRCA, CESC, CHOL, COAD, DLBC, ESCA, GBM, and HNSC(Fig 5C) (the ten tumor sets contain 60% samples of the TCGA data set) (we also analyzed the contribution of features on the remaining 23 cancer data sets to the results, see Fig S5). It can be observed from Fig S5 that the importance of mutation characteristics still accounts for a large proportion (from 25.16% to 66.90%). Meanwhile, we rank the Gini scores of the multi-omics features on the Pan-cancer data set and the 33 cancer types and select the ten most important features on each type of tumor (see Table S5 for details). The contribution of different omics data to different cancer types is different: the importance of DNA methylation data in UCEC, SKCM, and PAAD is higher than that in other cancer types, reaching 8.44%, 8.91%, and 11.90%, respectively.

## Discussion

Driver gene discovery is one of the critical issues in cancer genomics, which is critical for understanding cancer’s pathogenesis and advancing the development of targeted drugs. Inspired by the success of the transformers network in various fields of biomedicine, we propose a novel deep learning-based approach, Trans-Driver, which is the first to apply the transformer network in identifying cancer drivers. Our approach has significantly improved performance compared with other state-of-the-art driver gene discovery approaches and pinpointed a reasonable number of candidate driver genes by integrating four types of omics data across 33 TCGA tumor types. Also, Trans-Driver identified 185 alternative driver genes on the Pan-cancer set, of which 103 genes (55%) were included in the gold standard CGC data set, and 97 genes (51%) were included in the latest released driver gene set of TCGA. Trans-Driver reported 563 novel drivers across 33 cancer types based on the enrichment analysis. We found considerable evidence on Pan-cancer and individual tumor sets to support the novel cancer drivers identified by Trans-Driver through the literature search. For example, the candidate driver gene TFPI-2 is associated with BLCA [56], the candidate driver gene ELAVL1 is associated with BRCA cancer type [54], and the candidate driver gene PMG5 is associated with COAD cancer type [55]. Finally, we explained the association between features and model results and found that integrating multi-omics data could improve our approach’s performance compared to using only mutation data. Moreover, we found that the contribution of different omics data to the driver gene discovery varied on different tumor types.

Trans-Driver can identify cancer drivers that are difficult to detect only using somatic mutation data. For example, the ELF3 gene on BLCA is challenging to identify with other methods. The BRCA1 gene on BRCA, which is also more challenging to pinpoint, is successfully reported by our approach. The omics data analysis in these two cancer types showed that the mutation data contribute to the final prediction and other omics data (such as CNV, DNA methylation, somatic mutations, and RNA-seq data) play a crucial role. Integrating multi-omics data can provide a more comprehensive understanding of the biological processes of cancer driver genes. The fact that multi-omics data can be applied to predict driver genes further suggests connections across the omics data that can lead to a deeper understanding of cancer occurrence and progression mechanisms.

The Transformer started to be applied to machine translation tasks. Due to its superior network structural design has also achieved great success in computer vision and is comparable to convolutional neural networks. Just recently, Transformers were introduced into the biomedical field with great success. Senior previously used Transformer-based models to set a new height in protein 3D structure prediction at the CASP14 competition [57]. Protein will form a three-dimensional structure by crimping and folding, and the function of protein is determined by its structure. Understanding protein structure can help understand the role of proteins in the body and help develop drugs to treat diseases. Meanwhile, Avsec proposed Enformer, which uses an attention mechanism to process a more extensive range of DNA contextual information, dramatically improving predicting gene expression based on DNA sequences [58]. For the cancer driver gene discovery task, since its input is utterly different from Protein structure prediction and predicted gene expression, the introduction of Transformer requires a unique design combined with the corresponding model input. The combination of MLP, Transformer, and the imbalanced learning loss enables nonlinear modeling of low-dimensional correlated multi-omics data. It focuses more on the essential features, reducing the impact of unbalanced drivers and passengers on performance stability. The improvement of Trans-Driver compared with the DNN model shows the reasonableness of our novel framework.

Our framework has limitations. First, in the data preprocessing stage, Trans-Driver uses the mean value of all samples’ omics data to fill in the missing values without considering the correlation between different types of omics data. In the future work, we plan to use the method based on the generative adversarial network to learn the relationship between multi-omics data to improve the accuracy of Trans-Driver in cancer types with severe data deletion. Second, Trans-Driver used Pan-cancer data for model training, and due to the relative lack of samples on each cancer type, the model was not retrained. The model trained on the Pan-cancer data set was directly used for driver gene prediction on 33 cancer types. In the future, we expect to add the transfer learning technique to improve the accuracy of driver gene prediction on 33 cancer types. Third, the model architecture of Trans-Driver relies on deep learning, which requires higher quality omics data and its annotation than traditional statistical models. In the future, with the accumulation of more sample omics data on more cancer types, the trained models using Trans-Driver will be more accurate in predicting driver genes. Meanwhile, with the disclosure of genome-wide genomic data, we expect Trans-Driver to reveal the pathogenic mechanism of non-coding region driver genes and thus provide a deeper understanding of cancer occurrence and development.

## Acknowledgments

This work is supported by Natural Science Foundation of China under Grant No. 61902126, Shanghai Science and Technology Program “Distributed and generative fewshot algorithm and theory research” under Grant No. 20511100600, Shanghai Science and Technology Program “Federated based cross-domain and cross-task incremental learning” under Grant No. 21511100800.

## Author contributions

ZW designed the study. HY developed the Trans-Driver framework and implemented it. DL, JZ, and DZ collated ATAC-seq data and RNA-seq data from the TCGA database and completed the comparison. HY and LZ analyzed the experimental results. HY, LZ, and ZW wrote the manuscript. All authors read and approved the manuscript.

